# Classification of Healthy People and Schizophrenics Using Time-Frequency Domain Features Extracted from Electroencephalogram Signals

**DOI:** 10.64898/2026.04.13.718103

**Authors:** Nazila Ahmadi Daryakenari, Seyed Kamaleddin Setaredan

## Abstract

Schizophrenia (SZ) is a chronic and complex mental disorder associated with neurobiological deficits. The complexity and heterogeneity of schizophrenia symptoms pose challenges for objective diagnosis, which is currently based on behavioral and clinical manifestations. Furthermore, other psychiatric disorders such as bipolar disorder or major depressive disorder are often misdiagnosed as schizophrenia. Consequently, manual screening through psychiatrist-patient interviews is not entirely reliable. This study aims to develop an automated SZ diagnosis scheme using electroencephalogram (EEG) signals as a complementary tool to assist psychiatrists. A novel method is proposed, utilizing features from time, frequency, and time-frequency domains to classify EEG signals from schizophrenia patients and healthy individuals. Time-domain features, frequency-domain features, as well as nonlinear and statistical features were extracted, and 10 feature combinations were selected based on importance using a hybrid mutual information and Sequential Forward Feature Selection approach. Classification was performed using K-nearest neighbors (KNN), weighted KNN, linear and nonlinear support vector machines (SVM) with radial basis function kernels, decision trees, linear discriminant analysis, and the naive Bayes method. Remarkably, three classifiers achieved 100% accuracy.

## Introduction

Brain dysfunction due to disease or disorder significantly impacts normal human activity. Schizophrenia is a severe and chronic psychotic disorder characterized by a distortion in the perception of reality. The psychosis associated with schizophrenia involves symptoms such as delusions (persistent false beliefs), hallucinations (experiences of seeing, hearing, tasting, smelling, or feeling things that are not actually present, often auditory hallucinations like hearing voices in schizophrenia), disorganized speech and thinking, and abnormal motor behavior (ranging from childlike silliness to unpredictable or purposeless movements). Additional symptoms include social withdrawal, reduced emotional expression, and apathy [1].

Globally, schizophrenia is one of the 15 most disabling diseases. According to reports from the World Health Organization (WHO), approximately 26 million people worldwide suffer from schizophrenia. The cost of treatment is estimated to range from $32.5 billion to $65 billion annually, imposing a substantial economic burden. Moreover, nearly 5% of individuals with this disorder die by suicide. Schizophrenia typically begins in late adolescence or early adulthood, with symptoms often emerging gradually during this period.

Schizophrenia manifests earlier in men than in women and is more prevalent in urban settings and among minority groups [2]. However, the World Health Organization (WHO) has stated that this condition is treatable, and early diagnosis or diagnosis at later stages can be useful for determining the severity and phase of the disorder.

Schizophrenia imposes significant harm on patients, making timely and accurate diagnosis essential. Approximately 90% of untreated individuals with schizophrenia live in low- and middle-income countries. Traditional diagnosis relies on interviews with the patient conducted by a skilled psychiatrist, a process that is time-consuming, costly, exhausting, and prone to human error. Consequently, there is a need for a fully automated, relatively accurate, and cost-effective system for diagnosing schizophrenia, which can improve the efficiency of the current diagnostic process and serve as a supplementary tool [3].

Electroencephalography (EEG) is widely used for assessing and diagnosing brain functions and disorders due to its high temporal resolution and affordability. EEG signals are extensively used for schizophrenia diagnosis because they reflect the condition of the brain. This non-invasive technique has advantages over other imaging techniques, such as functional magnetic resonance imaging (fMRI), including greater flexibility in design, lower costs, and superior temporal resolution. EEG-based schizophrenia diagnosis plays a crucial role in early and precise detection, serving as a complementary tool in clinical practice [4].

Numerous studies have focused on classifying healthy individuals and schizophrenic patients [5] [6] [7] [8]. However, most research has been confined to a single domain, primarily the time or frequency domain.

In the proposed method, 18 feature models encompassing time-domain, frequency-domain, complexity, entropy, and statistical features were extracted from three domains: time, frequency, and time-frequency. A feature vector was created by integrating these domains. To reduce the dimensionality of the feature vector and select effective features, irrelevant features were eliminated using a mutual information method, and the optimal feature combination was selected using a Sequential Forward Feature Selection based on maximum performance (highest accuracy). Finally, classification was performed on the selected features using machine learning methods.

## Dataset and Methods

This section outlines the dataset used in this study, followed by discussions on preprocessing, feature extraction methods, feature selection, and classification.

### 1.1 Dataset

The dataset used in this study was recorded at the Atieh Clinical neuroscience center. It includes 14 healthy individuals and 14 individuals diagnosed with schizophrenia, with mean ages of 25.3±8.1 and 28.42±5.93 years, respectively. The sampling frequency was set at 500 Hz, and 19 electrodes were used following the international 10-20 system. Data were recorded in a resting state with eyes closed for 5-7 minutes, but only 5 minutes of data were used for uniformity and better comparability.

### 1.2 Preprocessing

Since brain signals have low power at frequencies above 50 Hz, and to eliminate power line interference, a finite impulse response (FIR) band-pass filter was used to filter out frequency components outside the range of 0.5 45 Hz. Additionally, as artifacts such as eye movement, motion noise, and heartbeat noise are not fully removed by band-pass filtering, independent component analysis (ICA) was employed to eliminate components that were not brain-related.

### 1.3 Feature Extraction

Features were extracted from three domains: time, frequency, and time-frequency.

- **Time Domain Features**:

Five important and widely used features were extracted:

- Zero Crossing Rate (ZCR)
- Hjorth Mobility
- Hjorth Complexity
- Higuchi’s Fractal Dimension
- Lempel-Ziv Complexity
- **Frequency Domain Features**:
- Mean power in delta, theta, alpha, beta, and gamma bands
- Delta-to-alpha power ratio
- Spectral entropy

- **Time-Frequency Domain Features**:

Discrete Wavelet Transform (DWT) was applied to extract features including:

- Kurtosis
- Hurst Exponent
- Shannon Entropy
- Log Energy Entropy
- Rényi Entropy
- Tsallis Entropy

The mother wavelet was selected based on similarity to the original signal. After testing various wavelets, Daubechies order 6 provided the best results and was employed in this study.

For time-domain features, calculations were performed once over the entire signal and then over specific frequency bands. In total, 18 feature models were used for processing.

#### Zero Crossing Rate (ZCR)

The Zero Crossing Rate (ZCR) represents the number of times a waveform crosses the x-axis. It is an indicator of signal noisiness and reflects signal complexity. High-frequency processes exhibit a high ZCR, while low-frequency processes show a slower ZCR.

Let *x*_*i*_*(n)*=0,1,…,*N*−1, represent the samples of the i-th window of the signal. The ZCR for the i-th window is computed as:

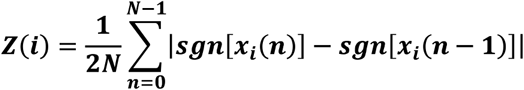

Here, N is the total number of samples in the window, and sgn is the sign function.

The average ZCR is calculated across all windows by first computing the ZCR for each window and then averaging these values to obtain the final ZCR[9].

#### Hjorth Mobility

Hjorth parameters are statistical features used in time-domain signal processing. Mobility is one of these parameters and represents the mean frequency of the signal. It provides information about the speed of the physiological process.

The mobility parameter, M, is calculated using the formula:

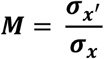

Where:

- σ X ′is the standard deviation of the first derivative of the signal.
- σX is the standard deviation of the original signal[10].

#### Hjorth Complexity

Hjorth Complexity quantifies the rate of frequency changes in a signal and describes its shape. It measures how similar a signal is to a pure sinusoidal wave and quantifies any deviation from this shape as an increase above unity.

The complexity parameter, C, is computed as:

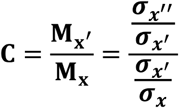

Where:

σx ″ is stand deviation of the second derivative of the signal

σ x is the standard deviation of the first derivative of the signal.

σx is the standard deviation of the original signal[10].

#### Higuchi Fractal Dimension

The Higuchi Fractal Dimension is a nonlinear measure of signal complexity in the time domain. It has been shown that the Higuchi method is a suitable numerical approach for quickly assessing the nonlinearity of a signal. Several studies analyzing EEG signals have demonstrated the superiority of the Higuchi method compared to Katz and Petrosian methods.

In the Higuchi method, the fractal dimension is obtained through the following steps:

Let X [1], X [2] X[3], …, X [N] be the time series under analysis, where N is the length of the original time series. A new time series 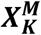 is created based on the following relationship:

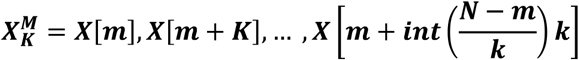

Where:

- M is the initial time value of 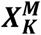,
- K is the time step.

The length of the new time series 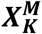 can be calculated using the following formula:

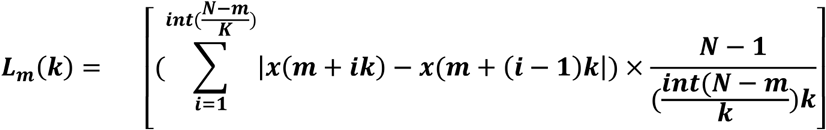

Where 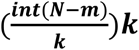 is a normalization factor. The overall mean length for all time series and for all k values, ranging from 1 to 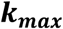, is given by:

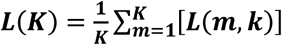

Finally, the slope of the best-fit linear function through the points 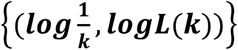 is defined as the Higuchi Fractal Dimension of the time series X [11].

#### Lempel-Ziv Complexity (LZC)

Lempel-Ziv Complexity (LZC) is a nonlinear analysis method used for data with short lengths. It is employed to assess the repetitiveness of EEG signals. This complexity measure is related to the number of distinct subsequences (or patterns) and their occurrence rates along a given sequence.

To compute LZC, the signal is first compared with a threshold and converted into a finite sequence of symbols. In medical signal analysis, the discrete medical signal is transformed into a binary sequence. Typically, the median is used as the threshold due to its resistance to outliers.

To calculate the LZ complexity, the sequence P is scanned from left to right. Each time a new subsequence of consecutive characters is encountered, the complexity counter c(n) is incremented[12].

#### Mean Band Power in Frequency Bands

One of the most common methods for analyzing EEG data is decomposing the signal into distinct frequency bands, such as delta (0.5-4 Hz), theta (4-8 Hz), alpha (8-12 Hz), beta (12-30 Hz), and gamma (30-100 Hz). All frequency bands are affected in schizophrenia. Mean band power involves calculating a single value that summarizes the contribution of the given frequency band.

To compute the mean band power, an estimate of the power spectral density must first be calculated. The power spectrum of a time series describes the distribution of power across its frequency components. The most commonly used method for this is the Welch periodogram, which involves averaging consecutive Fourier transforms of small overlapping or non-overlapping windows of the signal.

The absolute power of a band is the integral of all power values within that frequency range. Since no closed-form formula exists for integrating the areas under the absolute power curve of a specific frequency band, an approximation is used, typically employing Simpson’s composite rule. The area is divided into several rectangles, and then the areas of these rectangles are summed[13].

#### Power Ratio Between Two Frequency Bands

A common analysis metric in neurophysiological electrophysiology is calculating the power ratio between two frequency bands. Band ratio metrics are typically interpreted as reflecting the quantitative activity of oscillatory or periodic components, implicitly assuming that a ratio measures the relative power of two distinct oscillatory components that are well captured by predefined frequency bands.

An increase in delta power and a decrease in alpha power are observed in schizophrenia-related psychosis. Therefore, the delta-to-alpha power ratio has been studied, as this frequency ratio acts as an effective neuro-physiological biomarker[14].

#### Spectral Entropy

Spectral Entropy (SE) is a measure of the uncertainty or randomness in the pattern of a signal. It is essentially equivalent to the amount of information present in the signal. Spectral entropy quantifies the distribution of power across frequencies and indicates the uniformity of power distribution in the frequency domain.

The power spectral density P(w) represents the power spectrum of each frequency component of the signal, typically computed using the Welch method. Once the power spectral density is obtained, it is normalized, and spectral entropy can be calculated using Shannon’s entropy principle, as follows:

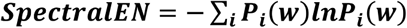

Where ***P***_***i***_ **(W)** is the power at the i-th frequency bin[15].

#### Kurtosis

In statistics and probability theory, Kurtosis describes the “peakedness” or flatness of a probability distribution. The more peak-like the shape of the probability density function, and the broader the tails or distribution, the higher the kurtosis value. Kurtosis is calculated as the normalized fourth moment[16].

#### Hurst Exponent

The Hurst Exponent (H) measures the long-term memory of time series data. It quantifies the tendency of a time series to revert to the mean or cluster in one direction. The Hurst exponent indicates the fractality of the signal and serves as a measure for signal variation. It ranges from 0 to 1, with higher values indicating smoother trends and lower volatility.

The Hurst exponent is defined by the following relation:

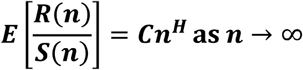

Where:

- R(n) is the cumulative range of the first n deviations from the mean,
- S(n) is the cumulative sum of the first n standard deviations,
- E[x] is the expected value of x,
- C is a constant, and
- n is the observation window size[17].

#### Shannon Entropy

Shannon Entropy measures the uncertainty of a random process. It quantifies the randomness in a signal and is interpreted as the impurity or unpredictability of the signal. Shannon entropy is calculated using the following formula:

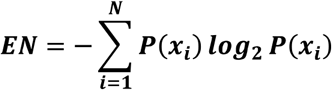

Where ***P* (*x***_***i***_) is the probability of occurrence of the event **x** _**i**_ [18].

#### Log Energy Entropy

Similar to wavelet entropy, Log Energy Entropy is used to calculate entropy based on the logarithmic energy of the signal. It can be computed using the following formula:

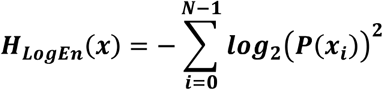

Where ***P (x***_***i***_) represents the probability of occurrence of the i-th event[18].

#### Rényi Entropy

In information theory, Rényi Entropy generalizes various forms of entropy, including Shannon entropy, Hartley entropy, and minimum entropy. It can be expressed as:

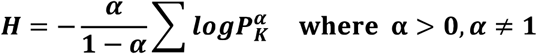

This entropy expression quantifies how likely it is that two independent random variables will take the same value, with α=2 corresponding to collision entropy [19].

#### Tsallis Entropy

Tsallis Entropy is a generalization of the Boltzmann-Gibbs entropy from statistical thermodynamics and is defined as:

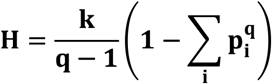

Where:

- k is a positive constant, and
- q is a non-extensive parameter.

For q>1, Tsallis entropy reacts more to frequent events, while for 0<q<1, it responds more strongly to rare events[18].

### 1.4 Feature Selection

Feature selection was carried out using a hybrid method combining mutual information and Sequential Forward Feature Selection. Initially, irrelevant features were removed using the scalar mutual information method, followed by the selection of the best feature combination based on the highest accuracy using the forward sequential search method. The advantage of the scalar method is that it has a low execution time, making it suitable for feature elimination, but it does not consider the relationship between features and classifier performance. Therefore, after this step, the Sequential Forward Feature Selection method was used to select the optimal feature combination.

#### Mutual Information

In probability theory and information theory, mutual information between two random variables is a measure of the mutual dependence between them. In other words, it quantifies how much information can be obtained about one random variable by observing the other. The concept of mutual information is inherently linked to the entropy of a random variable, which indicates the amount of information contained in a random variable. This method falls under scalar methods. Mutual information measures the similarity between the joint distribution p(X,Y) and the product of the marginal probabilities p(X)p(Y).

#### Sequential Forward Feature Selection

The Sequential Forward Feature Selection method belongs to vector-based feature selection techniques. The process begins by selecting the best single feature with the highest performance (based on the highest accuracy). Then, a feature pair is formed using one of the remaining features and the best feature, and the best pair is selected. In the next step, a triplet of features is formed using the two best features and another remaining feature, and the best triplet is chosen. This process continues until the specified number of features is selected. The steps are as follows:

1. Start with an empty set: **Y**_**0**_**={Ø}**
2. The best feature is selected using the following criterion:

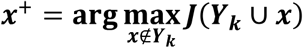
3. The set is updated:

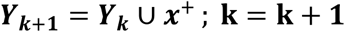
4. Return to step 2[19].

### 1.5 Classification of Features

#### K-Nearest Neighbors (KNN)

The K-Nearest Neighbors (KNN) algorithm is one of the simplest and oldest classification algorithms. KNN is a supervised learning technique that can be considered a simplified version of the Bayesian classifier. In the KNN algorithm, K represents the number of nearest neighbors considered for simple majority voting[20].

#### Weighted K-Nearest Neighbors (WKNN)

Similar to the K-Nearest Neighbors algorithm, but in Weighted K-Nearest Neighbors (WKNN), each neighbor is given a weight based on its distance from the test sample. Neighbors that are closer to the test point have a higher weight, and those further away have a lower weight when voting. The distance between neighbors is typically calculated using Euclidean distance[20].

#### Linear Support Vector Machine (SVM)

A Linear Support Vector Machine (SVM) is a supervised machine learning algorithm. This algorithm aims to find the decision boundary that maximizes the distance between data points from different classes. By doing so, it ensures that the classification decision is as accurate as possible, by creating the largest possible margin between the classes[20].

#### Nonlinear Support Vector Machine (SVM)

In the Nonlinear Support Vector Machine (SVM), the data is initially transformed using a nonlinear mapping function, such as the Radial Basis Function (RBF) kernel, from the original space to a higher-dimensional space. This transformation reduces the overlap between classes in the higher-dimensional space. After mapping the data to the higher-dimensional space, an optimal hyperplane is obtained that maximizes the margin between classes[20].

#### Decision Tree (DT)

Decision Tree (DT) is one of the earliest and most prominent machine learning algorithms. A decision tree models decision-making logic. It examines the data branch by branch, selecting the best feature at each branch based on the highest performance to separate the unsorted dataset. The data is then classified according to the selected feature[20].

#### Linear Discriminant Analysis (LDA)

Linear Discriminant Analysis (LDA) is a linear classifier. It finds a suitable line or hyperplane in an optimal direction that projects the data onto it, thereby reducing dimensionality. The line or hyperplane is chosen such that the within-class variance is minimized, and the between-class variance is maximized[20].

#### Naive Bayes (NB)

Naive Bayes (NB) is a classification technique based on Bayes’ theorem. It computes the probability of an event based on prior knowledge of the conditions related to that event. This classifier assumes that a specific feature within a class is independent of other features, though features within the class can be mutually dependent. Using Bayes’ theorem, the posterior probability of a feature vector is calculated. The test sample is assigned to the class that has the highest posterior probability[20].

### 1.6 Validation

To prevent overfitting and ensure more reliable results with a small dataset, 10-fold cross-validation and leave-one-out cross-validation methods were used.

In 10-fold cross-validation, the data is divided into 10 subsets. Each subset is randomly assigned, and in each iteration, 9 subsets are used for training, and the remaining one is used for testing. After completing all iterations, the accuracy values obtained are summed and averaged, with the reported accuracy being the average of these 10 values.

In leave-one-out cross-validation, all data except one sample is used for training, and the remaining sample is used for testing. This process is repeated N times, where N represents the number of data points. In this study, N=28, and the final accuracy is the average of the 28 values. The advantage of this method is that it uses all available data for both training and testing the model.

## Results

Since the vector-based methods introduce and select the best feature combinations separately according to each classifier’s perspective, the Sequential Forward Feature Selection was applied to the feature sets extracted from the three domains individually to choose the best feature combination for each classifier. To enhance processing speed, reduce the number of features, and prevent overfitting (given that each group contains only 14 individuals), the top 10 features that, when combined, yielded the highest performance (i.e., the highest accuracy) were selected for each classifier. The results for each classifier, based on Sequential Forward Feature Selection method, are shown in Figures (1) to (7). These figures display the number of feature combinations on the X-axis and the corresponding performance (accuracy) based on the feature combinations on the Y-axis.

**Figure 1:**
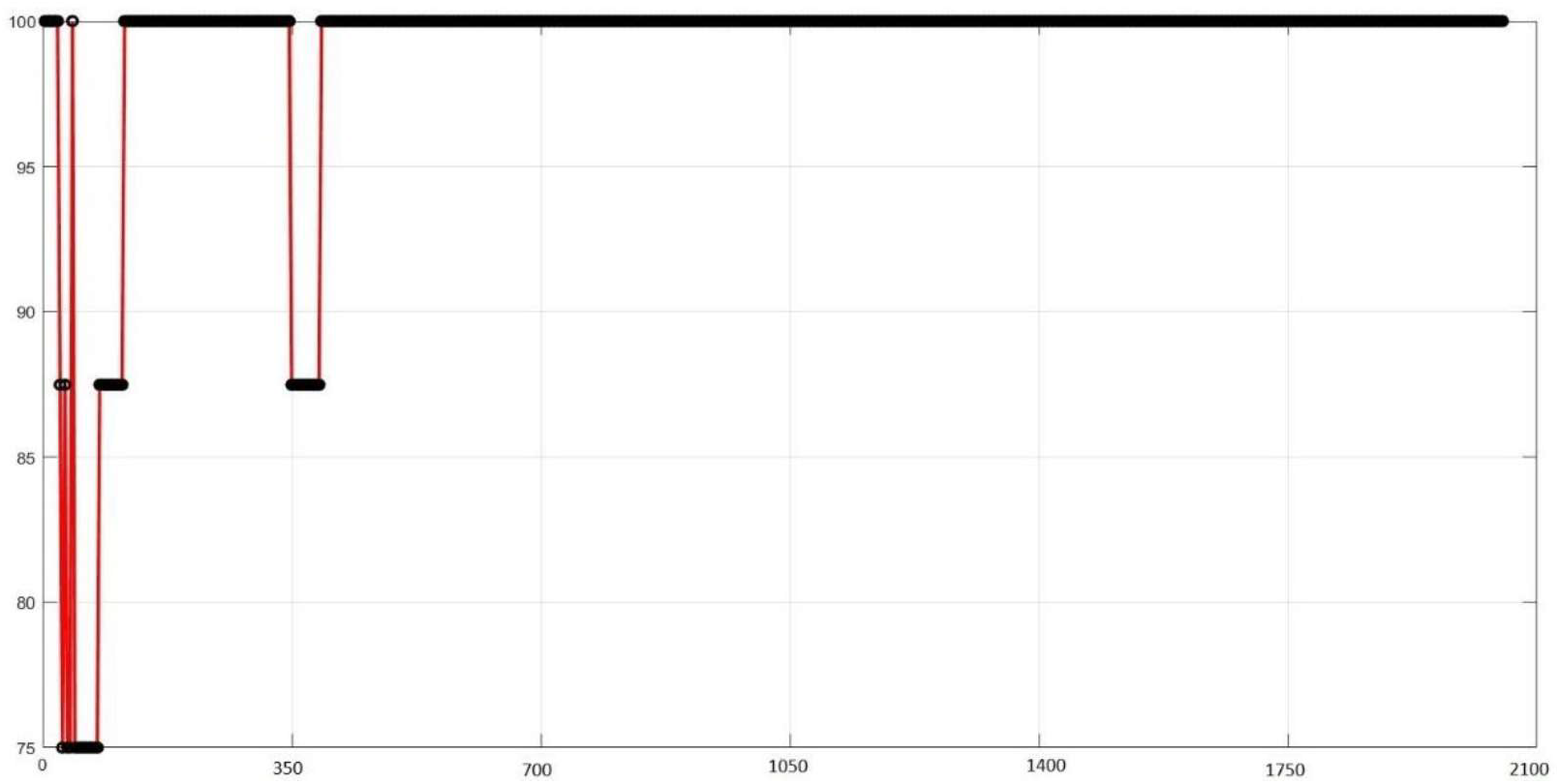
Results of the Sequential Forward Feature Selection Method for the KNN Classifier

**Figure 2:**
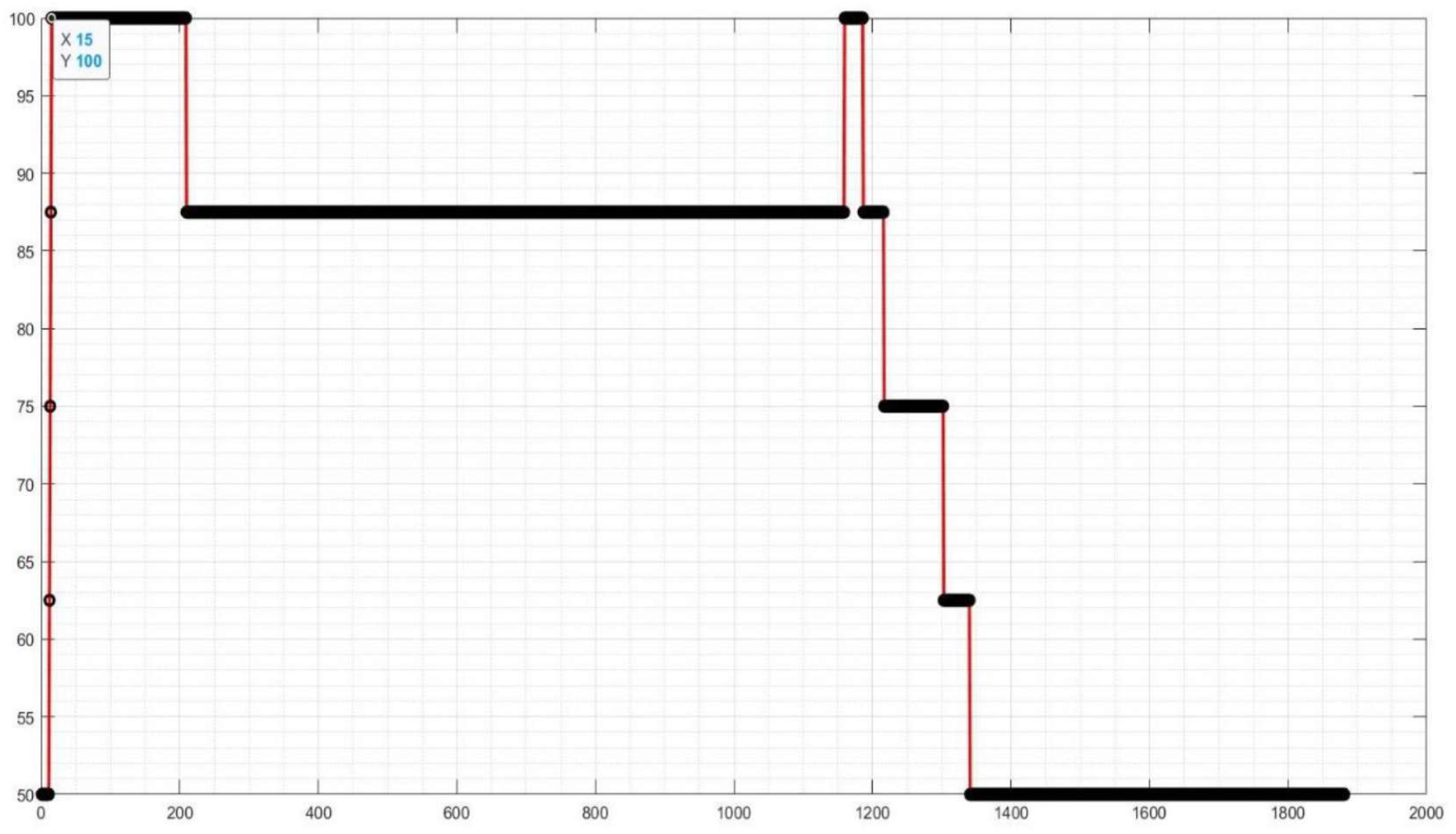
Results of the Sequential Forward Feature Selection Method for the WKNN Classifier

**Figure 3:**
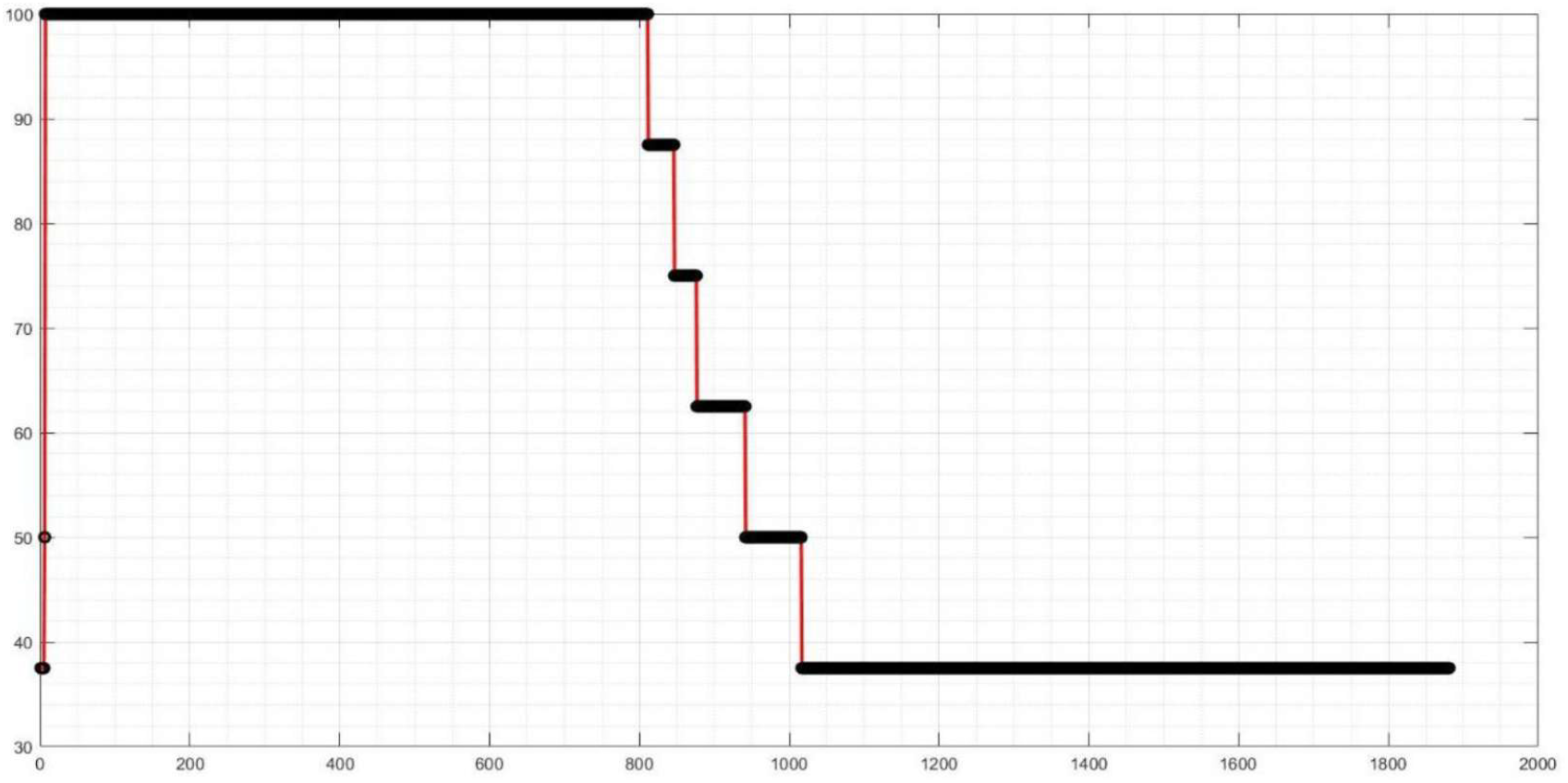
Results of the Sequential Forward Feature Selection Method for the Linear SVM Classifier

**Figure 4:**
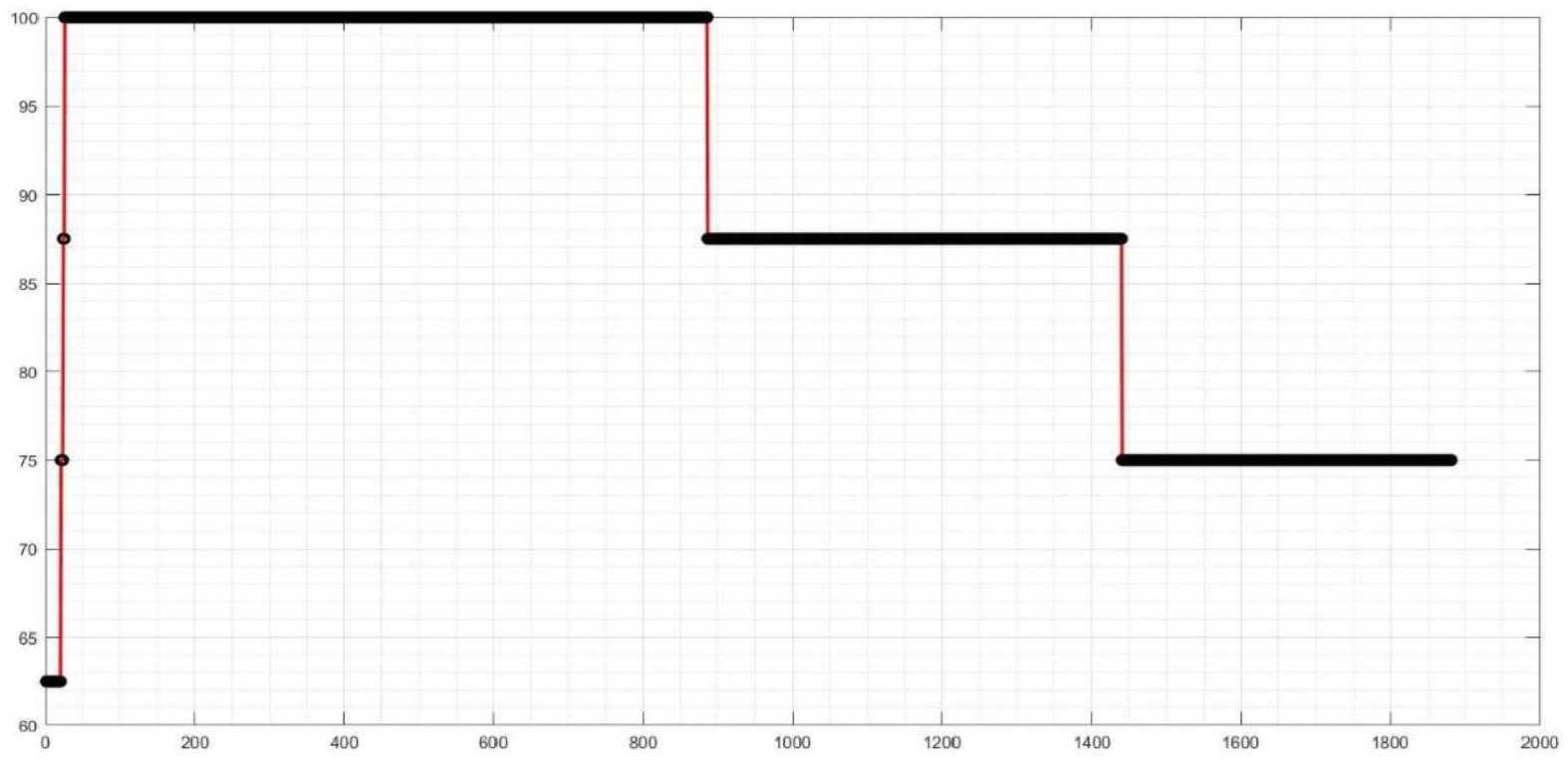
Results of the Sequential Forward Feature Selection Method for the non-Linear SVM Classifier

**Figure 5:**
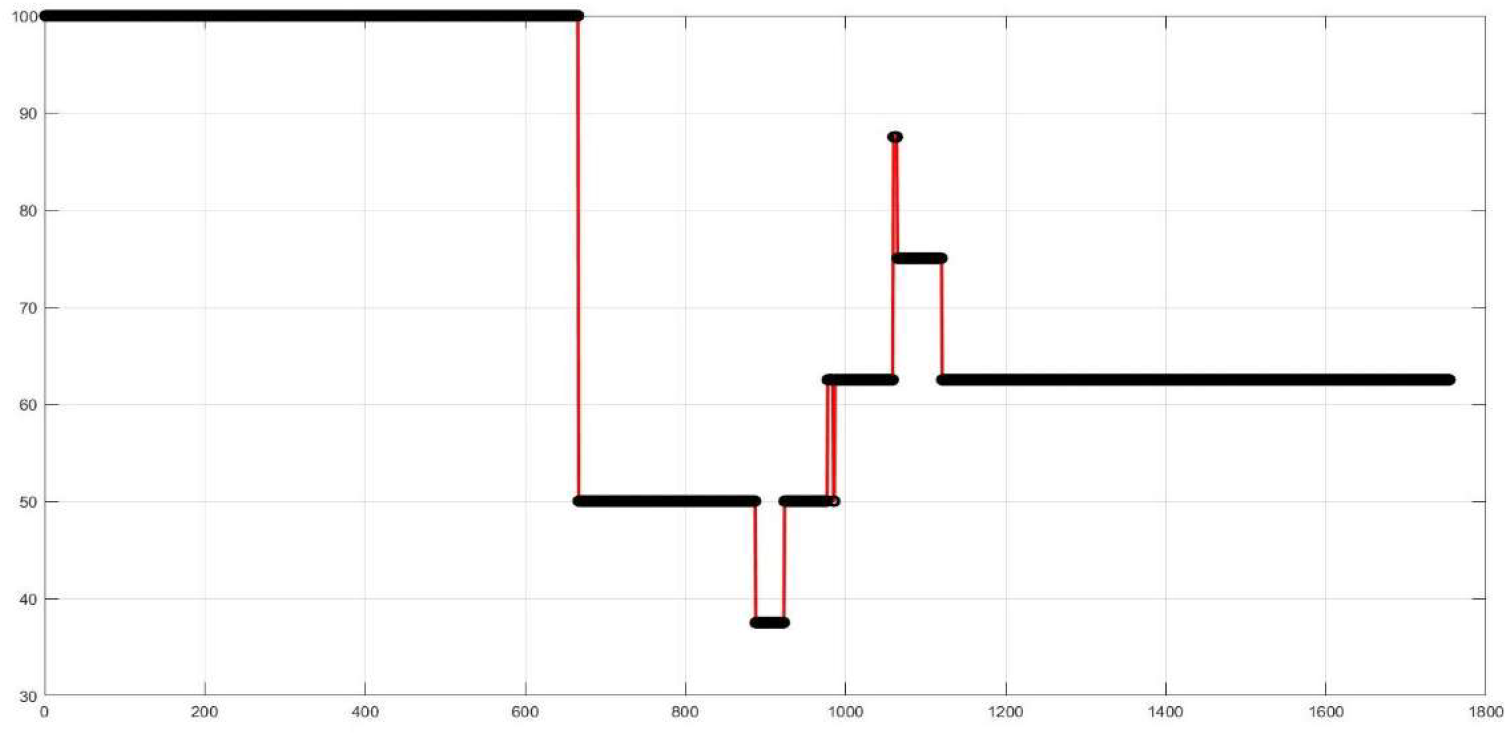
Results of the Sequential Forward Feature Selection Method for the LDA Classifier

**Figure 6:**
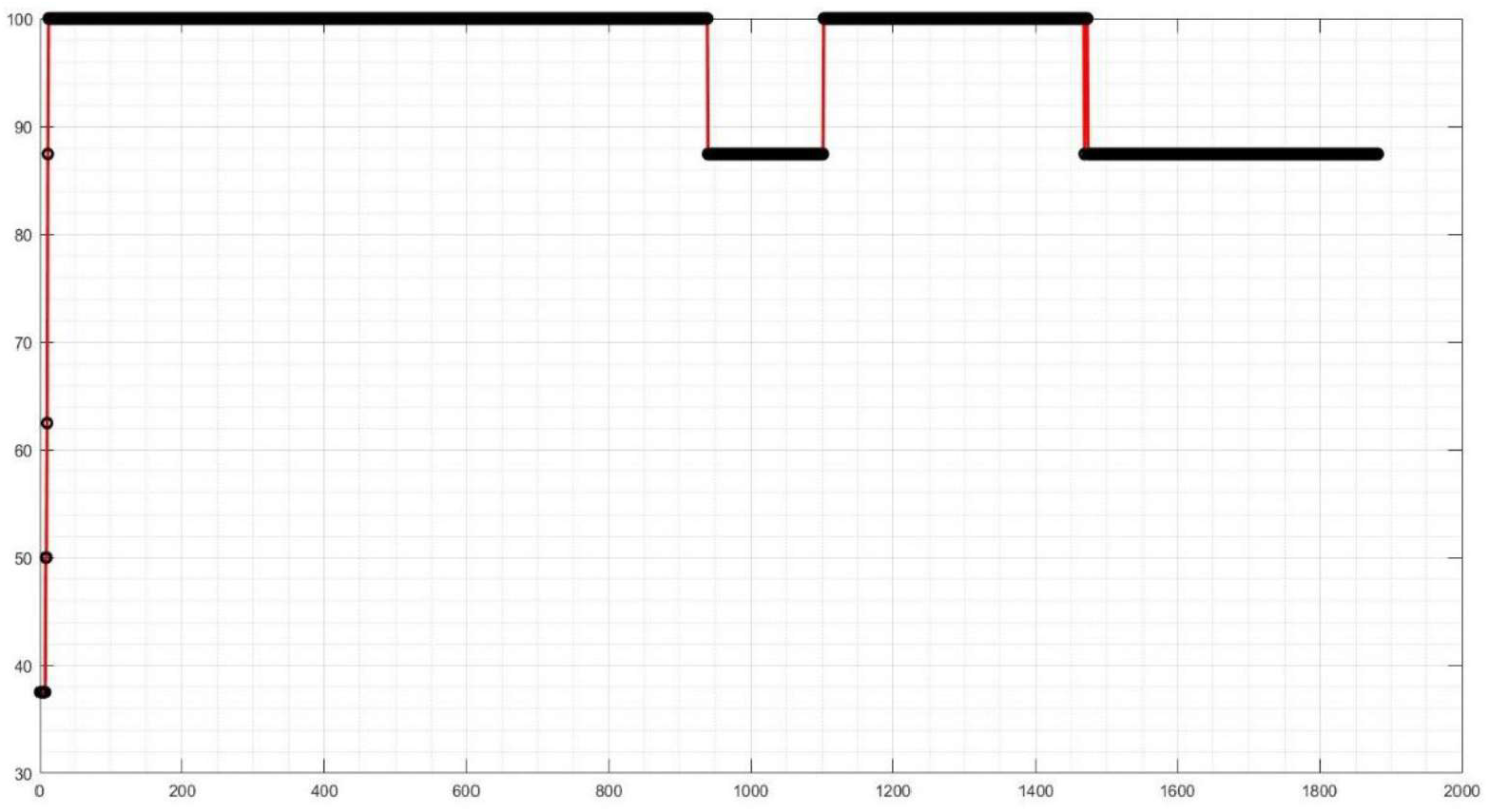
Results of the Sequential Forward Feature Selection Method for the Decision Tree Classifier

**Figure 7:**
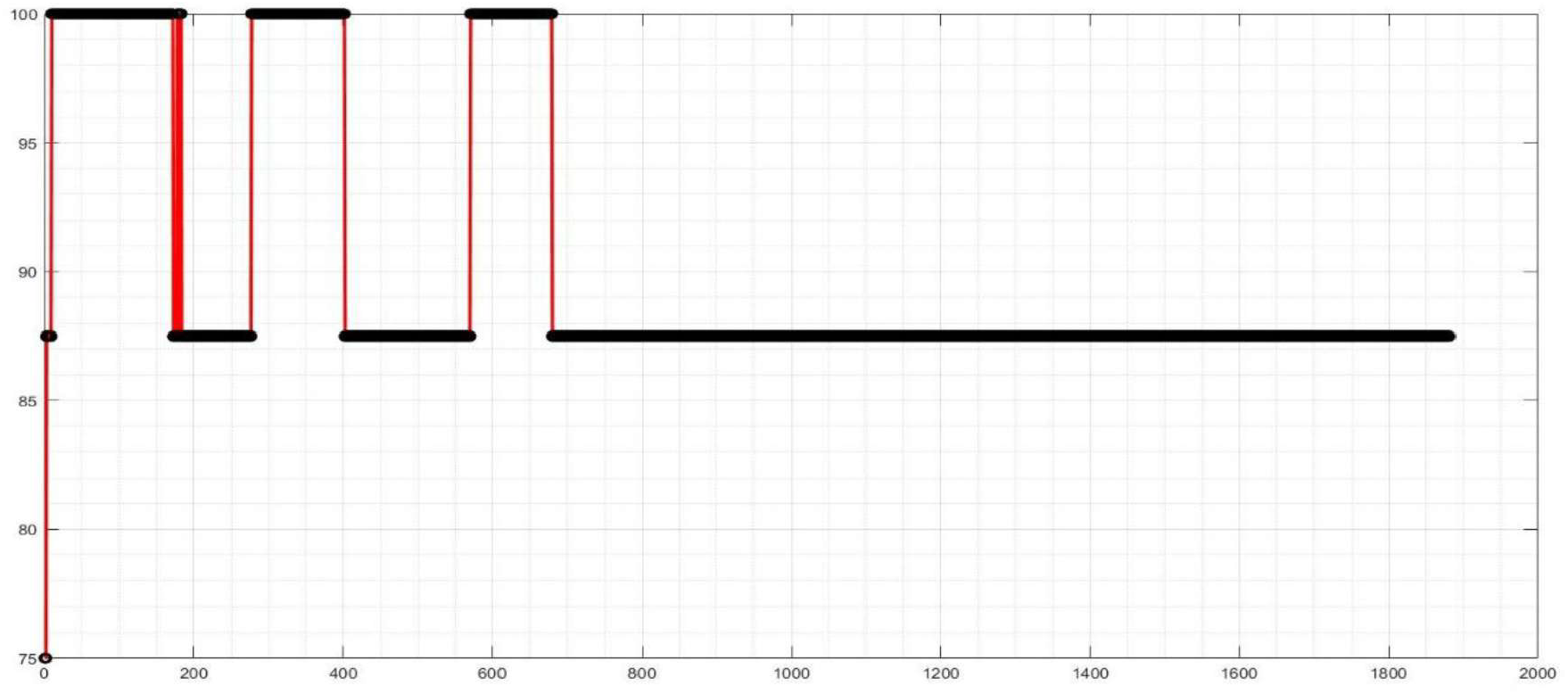
Results of the Sequential Forward Feature Selection Method for the Naive Bayes Classifier

This table lists the top 10 feature combinations selected for each classifier using the Sequential Forward Feature Selection method. The feature numbers correspond to the most effective combinations for achieving the highest accuracy in each classifier.

This table provides the types of features corresponding to the feature numbers listed in Table 1. Each selected feature type is associated with a feature number, showing the specific characteristics (e.g., complexity, power, entropy) chosen for each classifier.

**Table 1:** Selected Feature Numbers for Each Classifier Using the Forward Sequential Search Method.

| Classifier | Selected Feature Numbers |
| --- | --- |
| KNN | 5, 1, 3, 10, 17, 8, 6, 12, 13, 9 |
| WKNN | 5, 1, 2, 10, 18, 8, 6, 12, 14, 9 |
| Linear SVM | 5, 1, 2, 3, 6, 36, 7, 49, 21, 35 |
| Nonlinear SVM | 5, 1, 3, 4, 6, 36, 7, 45, 21, 30 |
| LDA | 5, 1, 2, 3, 4, 6, 7, 8, 10, 11 |
| Decision Tree | 5, 1, 2, 3, 4, 6, 7, 8, 9, 10 |
| Naive Bayes | 5, 1, 2, 3, 7, 8, 12, 13, 9, 14 |

**Table 2:**
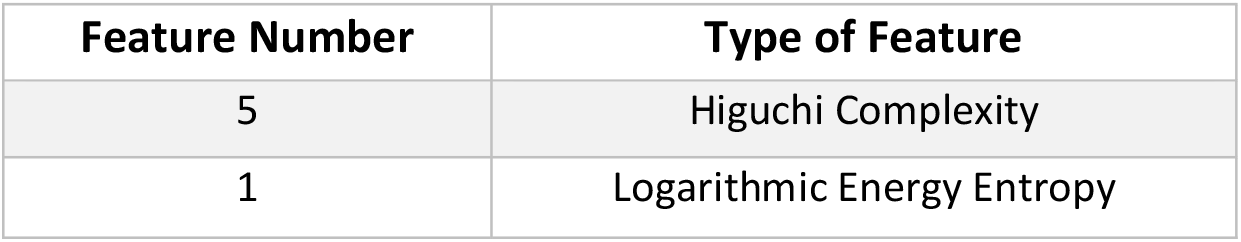
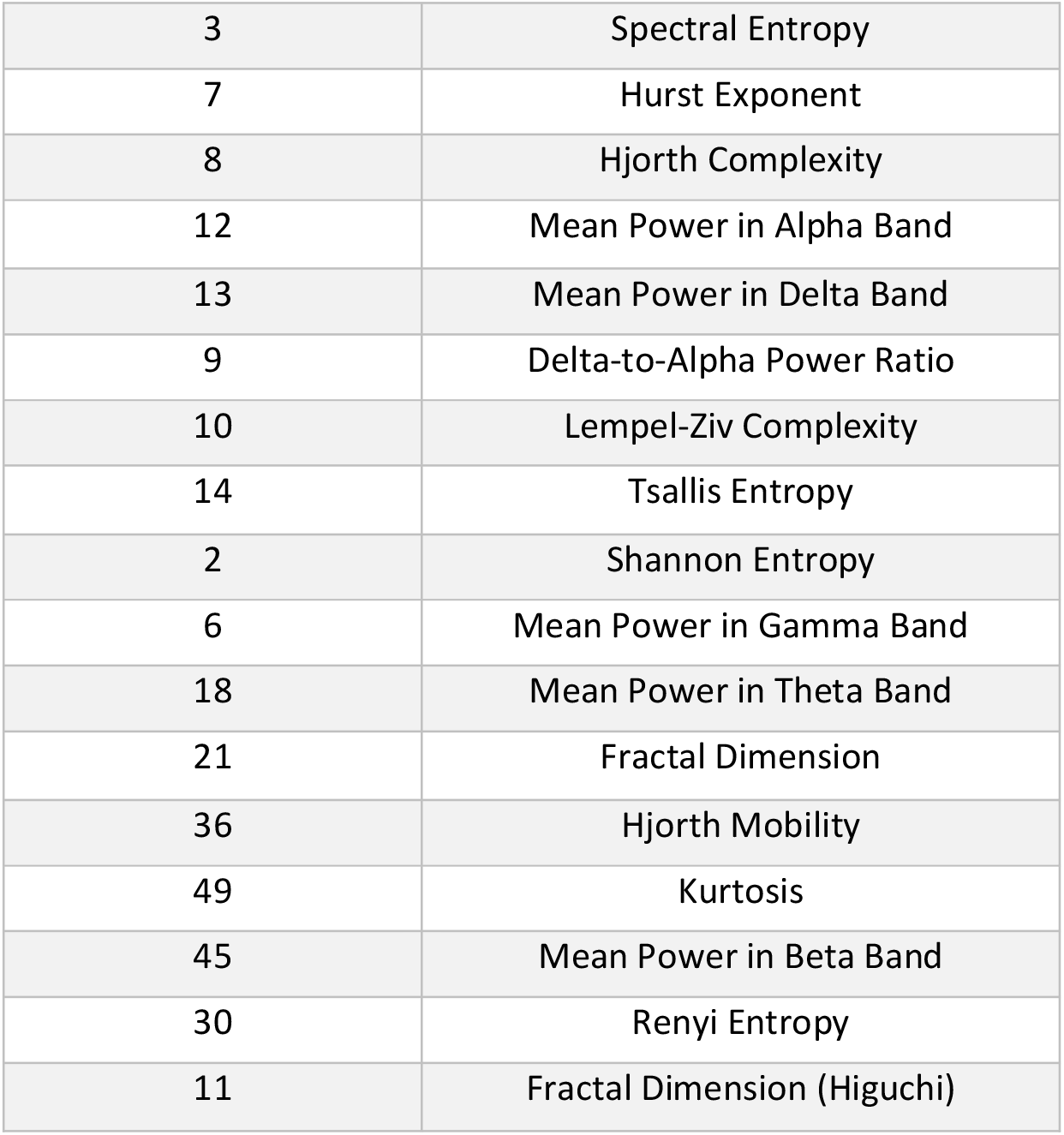
Type of Features Selected for Each Classifier Using the Sequential Forward Feature Selection Method.

As observed, Lempel-Ziv Complexity from the time domain, Spectral Entropy from the frequency domain, and Logarithmic Energy Entropy from the time-frequency domain were common across all classifiers and represented the best feature combinations based on performance.

The classification results for each domain (time, frequency, and time-frequency) after feature selection are provided in Tables 3 to 5. All results are based on the Leave-One-Out cross-validation method.

**Table 3:** Classification Results Using Leave-One-Out Validation for the Time Domain.

| Classifier | Specificity (%) | Sensitivity (%) | Accuracy (%) |
| --- | --- | --- | --- |
| KNN (k=3) | 90 | 100 | 95 |
| WKNN | 96 | 96 | 96 |
| Linear SVM | 92 | 100 | 96.42 |
| Nonlinear SVM | 92.85 | 100 | 96 |
| LDA | 92 | 100 | 96 |
| Decision Tree | 92.85 | 92.85 | 92.85 |
| Naive Bayes | 100 | 90 | 95 |

**Table 4:** Classification Results Using Leave-One-Out Validation for the Frequency Domain.

| Classifier | Specificity (%) | Sensitivity (%) | Accuracy (%) |
| --- | --- | --- | --- |
| KNN (k=3) | 80 | 80 | 80 |
| WKNN | 90 | 90 | 90 |
| Linear SVM | 90 | 90 | 90 |
| Nonlinear SVM | 100 | 80 | 90 |
| LDA | 100 | 100 | 100 |
| Decision Tree | 92.85 | 92.85 | 92.85 |
| Naive Bayes | 100 | 92.85 | 96.42 |

**Table 5:** Classification Results Using Leave-One-Out Validation for the Time-Frequency Domain.

| Classifier | Specificity (%) | Sensitivity (%) | Accuracy (%) |
| --- | --- | --- | --- |
| KNN (k=3) | 85.71 | 85.71 | 85.71 |
| WKNN | 92 | 92 | 92 |
| Linear SVM | 92 | 100 | 96 |
| Nonlinear SVM | 90 | 100 | 95 |
| LDA | 100 | 90 | 95 |
| Decision Tree | 92.85 | 92.85 | 92.85 |
| Naive Bayes | 100 | 100 | 100 |

These tables present the classification performance based on the Leave-One-Out cross-validation method for the three feature domains: time, frequency, and time-frequency. The accuracy values are consistent across classifiers for each domain and provide insights into the feature domains’ effectiveness.

As observed, the Naive Bayes classifier achieved the best performance, while the KNN classifier performed the worst.

In Tables 6 to 8, the classification results using 10-fold cross-validation and Leave-One-Out cross-validation are presented, both before and after feature selection, for the integration of the three domains (time, frequency, and time-frequency).

**Table 6:** Classification Results Using 10-Fold Cross-Validation Before Feature Selection for the Three Domains.

| Classifier | Specificity (%) | Sensitivity (%) | Accuracy (%) |
| --- | --- | --- | --- |
| KNN (k=3) | 90 | 90 | 90 |
| WKNN | 90 | 80 | 85 |
| Linear SVM | 90 | 100 | 95 |
| Nonlinear SVM | 90 | 100 | 95 |
| LDA | 90 | 100 | 95 |
| Decision Tree | 100 | 90 | 95 |
| Naive Bayes | 100 | 90 | 95 |

**Table 7:** Classification Results Using Leave-One-Out Cross-Validation Before Feature Selection for the Three Domains.

| Classifier | Specificity (%) | Sensitivity (%) | Accuracy (%) |
| --- | --- | --- | --- |
| KNN (k=3) | 85 | 85 | 85 |
| WKNN | 80 | 80 | 80 |
| Linear SVM | 90 | 100 | 95 |
| Nonlinear SVM | 100 | 90 | 95 |
| LDA | 92.85 | 92.85 | 92.85 |
| Decision Tree | 100 | 92.85 | 96.42 |
| Naive Bayes | 100 | 100 | 100 |

**Table 8:** Classification Results Using 10-Fold Cross-Validation After Feature Selection for the Three Domains.

| Classifier | Specificity (%) | Sensitivity (%) | Accuracy (%) |
| --- | --- | --- | --- |
| KNN (k=3) | 96 | 100 | 98 |
| WKNN | 90 | 100 | 95 |
| Linear SVM | 100 | 100 | 100 |
| Nonlinear SVM | 100 | 100 | 100 |
| LDA | 94 | 98 | 96 |
| Decision Tree | 100 | 100 | 100 |
| Naive Bayes | 98 | 98 | 98 |

**Table 9:**
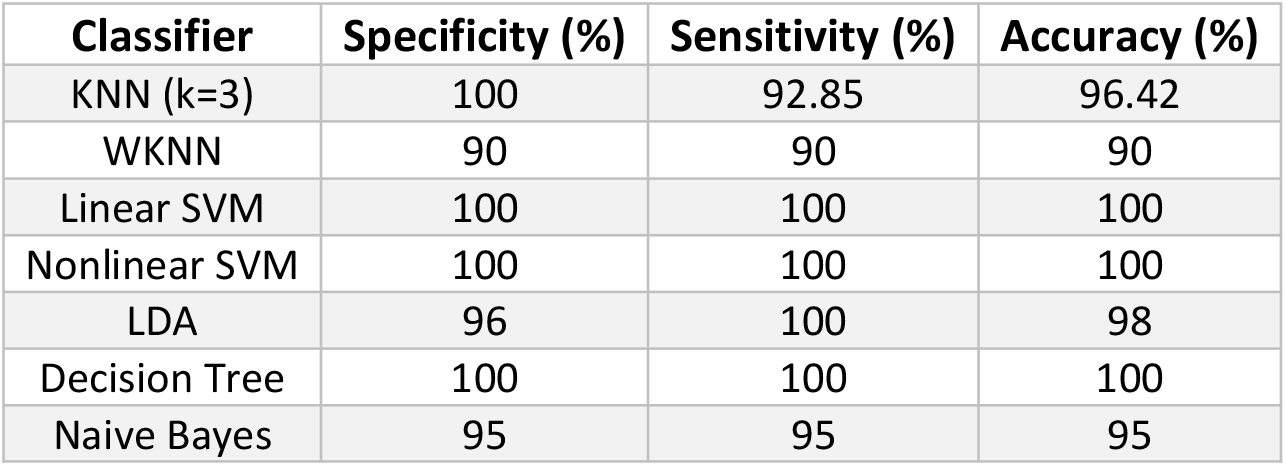
Classification Results Using Leave-One-Out Cross-Validation After Feature Selection for the Three Domains.

| Classifier | Specificity (%) | Sensitivity (%) | Accuracy (%) |
| --- | --- | --- | --- |
| KNN (k=3) | 100 | 92.85 | 96.42 |
| WKNN | 90 | 90 | 90 |
| Linear SVM | 100 | 100 | 100 |
| Nonlinear SVM | 100 | 100 | 100 |
| LDA | 96 | 100 | 98 |
| Decision Tree | 100 | 100 | 100 |
| Naive Bayes | 95 | 95 | 95 |

## Conclusion

The use of three domains for feature extraction and the hybrid method for feature selection has led to an accuracy of 100%, which can be utilized as a complementary tool for disease diagnosis. In future work, this method can be applied to other diseases, or for classification between positive and negative symptoms, or various types of schizophrenia. Furthermore, if a dataset specific to schizophrenia in adolescents becomes available, the selected features can be applied to both groups, and the results can be compared between schizophrenia in adolescents and adults.

